# 4-hydroxyphenylpyruvate dioxygenase thermolability is responsible for temperature-dependent melanogenesis in *Aeromonas salmonicida* subsp. *salmonicida*

**DOI:** 10.1101/324731

**Authors:** Yunqian Qiao, Jiao Wang, He Wang, Baozhong Chai, Chufeng Rao, Xiangdong Chen, Shishen Du

## Abstract

*Aeromonas salmonicida* subsp. *salmonicida* (*A.s.s*) is a major pathogen affecting fisheries worldwide. It is a well-known member of the pigmented *Aeromonas* species, which produces melanin at ≤ 22 °C. However, melanogenesis decreases as the culture temperature increases and is completely suppressed at 30-35 °C while bacterial growth is not affected. The mechanism and biological significance of this temperature-dependent melanogenesis are not clear. Heterologous expression of an *A.s.s.* 4-hydroxyphenylpyruvate dioxygenase (HppD), the most crucial enzyme in the HGA-melanin synthesis pathway, results in thermosensitive pigmentation in *Escherichia coli*, suggesting that HppD plays a key role in this process. In the current study, we demonstrated that the extreme thermolability of HppD is responsible for the temperature-dependent melanization of *A.s.s.* Substitutions in three residues, Ser18, Pro103, or Leu119 of HppD from *A.s.s* increases the thermolability of this enzyme and results in temperature-independent melanogenesis. Moreover, replacing the corresponding residues of HppD from *Aeromonas* media strain WS, which forms pigment independent of temperature, with those of *A.s.s* HppD leads to thermosensitive melanogenesis. Structural analysis suggested that mutations at these sites, especially at position P103, can strengthen the secondary structure of HppD and greatly improve its thermal stability. In addition, we found that HppD sequences of all *A.s.s* isolates are identical and that two of the three residues are completely conserved within *A.s.s* isolates, which clearly distinguishes these from other *Aeromonas* strains. We suggest that this property represents an adaptive strategy to the psychrophilic lifestyle of *A.s.s.*

**Importance:** *Aeromonas salmonicida* subsp. *salmonicida* (*A.s.s*) is the causative agent of furunculosis, a bacterial septicemia of cold water fish of the *Salmonidae* family. As it has a well-defined host range, *A.s.s* has become an ideal model to investigate the co-evolution of host and pathogen. For many pathogens, melanin production is associated with virulence. Although other species of *Aeromonas* can produce melanin, *A.s.s* is the only member of this genus that has been reported to exhibit temperature-dependent melanization. Here we demonstrate that thermosensitive melanogenesis in *A.s.s* strains is due to the thermolability of 4-hydroxyphenylpyruvate dioxygenase (HppD). The strictly conserved *hppD* sequences among *A.s.s* and the exclusive thermosensitive pigmentation of these strains might provide insight into the role of melanin in the adaptation to a particular host, and offer a novel molecular marker to readily differentiate *A.s.s* strains from other *A. salmonicida* subspecies and *Aeromonas* species.

## Introduction

Melanins are structurally complex high molecular weight pigments formed by the oxidative polymerization of phenolic and/or indolic compounds (1). They represent the most ubiquitous natural pigment and are produced by organisms from every kingdom (2). Melanin has various functions in diverse organisms, and it is believed that it confers a survival advantage for microorganisms, especially under stressful environmental conditions such as radiation, oxidative stress, the presence of digestive enzymes, and heavy metal toxicity (2). These molecules have also been associated with the virulence and pathogenicity of a variety of microbes such as *Cryptococcus neoformans*, *Wangiella dermatitidis*, and *Vibrio cholerae* (3–5), by protecting pathogens against host defenses and interfering with the host immune response (2). Melanins are classified into several categories based on their synthesis pathway, and bacteria usually synthesize two types, namely eumelanin or pyomelanin. Eumelanin is catalyzed by tyrosinase or other polyphenol oxidases to hydroxylate tyrosine, via the intermediate L-3,4-dihydroxyphenylalanine (L-DOPA), into dopaquinone, which then auto-oxidizes and polymerizes to form melanin (6). Pyomelanin is synthesized by the homogentisic acid (HGA) pathway, in which 4-hydroxyphenylpyruvate is converted to HGA by 4-hydroxyphenylpyruvate dioxygenase (HppD), which is followed by the oxidative polymerization of HGA to produce the pigment (6). Our previous study showed that *Aeromonas media* WS, a strain that produces melanin at high levels, mainly synthesizes pyomelanin through the HGA pathway (7).

The *Aeromonas* genus is a collection of gram-negative, rod-shaped, facultatively anaerobic bacteria that include motile, non-motile, mesophilic, and psychrophilic species. They are widely distributed in aquatic environments and have been frequently isolated from aquatic animal species. Some *Aeromonas* strains are principal or opportunistic pathogens of fish and other cold-blooded animals, as well as the causative agent of infectious complications in humans (8). Melanogenesis has been described in some species of this genus, such as *A. media*, *Aeromonas salmonicida*, *Aeromonas bestiarum*, and *Aeromonas eucrenophila* (9). However, a number of *Aeromonas* species are not considered to produce melanin, including *Aeromonas hydrophila*, *Aeromonas allosaccharophila*, and *Aeromonas encheleia* (10). In fact, melanin synthesis is one of the key criteria for the taxonomy of this genus.

Among melanogenic *Aeromonas* species, *A. salmonicida* subsp. *salmonicida* (*A.s.s*), an important salmonid fish pathogen and the etiologic agent of furunculosis, is the only member that has been described to exhibit temperature-dependent melanogenesis. It has been demonstrated that *A.s.s* can initiate pigmentation at lower temperatures such as 22 °C. However, this process is thermosensitive. The level of melanin production reduces as the incubation temperature increases, and this process is completely suppressed at temperatures > 30 °C, at which the bacteria still grow normally (11–13). Nevertheless, the reason for this phenomenon is not yet clear. Increasing the temperature can also alter some other characteristics of this bacterium, including the loss of virulence (> 25 °C) (14–16) and the auto-agglutination trait (>25 °C) (17). Since *A.s.s* is psychrophilic (optimal growth at 22-25 °C), these temperature-dependent phenotypes probably represent adaptations to psychrophilic environments.

In the recent study, we suggested that *Aeromonas* species mainly produce pyomelanin through the HGA pathway (7) and showed that heterologous expression of *hppD* from an *A.s.s* strain in *Escherichia coli* BL21 leads to similar temperature-dependent pigmentation. This result suggests that the abolished pigmentation of *A.s.s* at elevated temperatures is likely due to the thermolabile activity of HppD. In this study, we confirmed this hypothesis and identified the critical residues of HppD, specifically, S18, P103, and L119, that are responsible for its thermolability. Substitutions at these residues resulted in temperature-independent melanogenesis in *A.s.s.* Moreover, replacing the corresponding residues of HppD from other *Aeromonas* species with those from *A.s.s* resulted in thermal sensitive melanization. Taken together, our results demonstrate that the thermosensitive melanogenesis of *A.s.s* is dependent on individual residues in the HppD protein.

## Results

### *hppD* genes of *A.s.s* isolates are identical

To investigate the molecular mechanism responsible for the temperature-dependent pigmentation of *A.s.s*, we first ensured that this phenotype is ubiquitous in various *A.s.s* isolates. As shown in Fig. S1, all four *A.s.s* strains maintained in our laboratory (listed in Table 1; AB98041, KACC14791, 2013-8, and 2014-235) exhibited similar thermosensitive pigmentation as reported in the literature. We reasoned that if HppD actually plays an important role in this phenotype, the amino acid sequences of HppD from various *A.s.s* isolates should be highly conserved. Thus, we cloned and sequenced *hppD* genes from the four isolates and discovered that the sequences were 100% identical at the nucleotide level. Furthermore, alignment of the sequences of *hppD* from all 26 *A.s.s* isolates available in GenBank revealed that *hppD* was 100% conserved at the nucleotide level (data not shown), even though these strains were isolated in different years, from diverse areas, and from a variety of hosts (see detailed strain information in Table S1). Therefore, the thermal sensitive melanization observed in *A.s.s* isolates is likely due to the different activities of HppD at different temperatures. For simplicity, we refer to the *hppD* gene from *A.s.s* strains as *hppD*-AS in the rest of this paper.

**Table 1.**
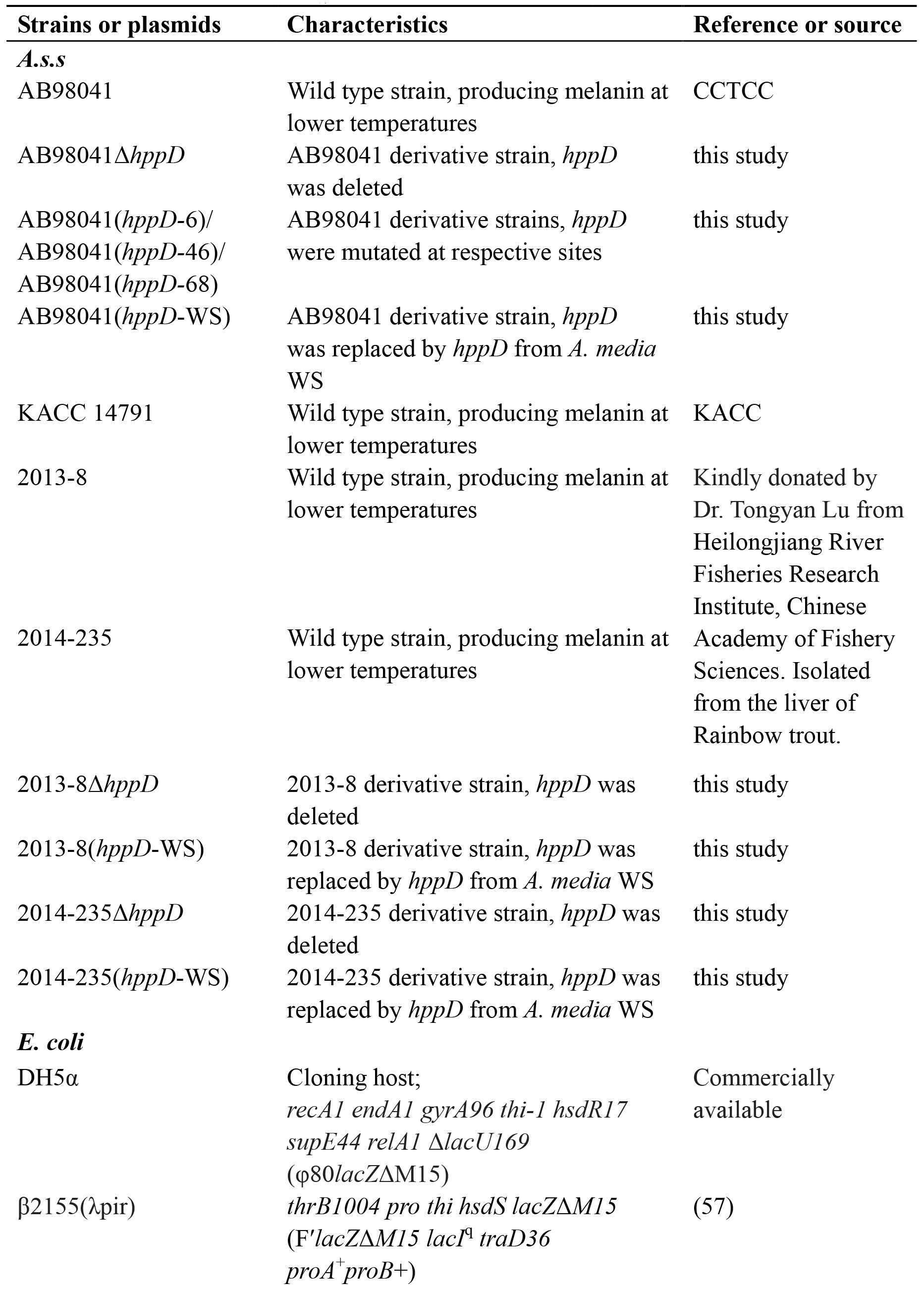
Bacterial strains and plasmids used in this study.

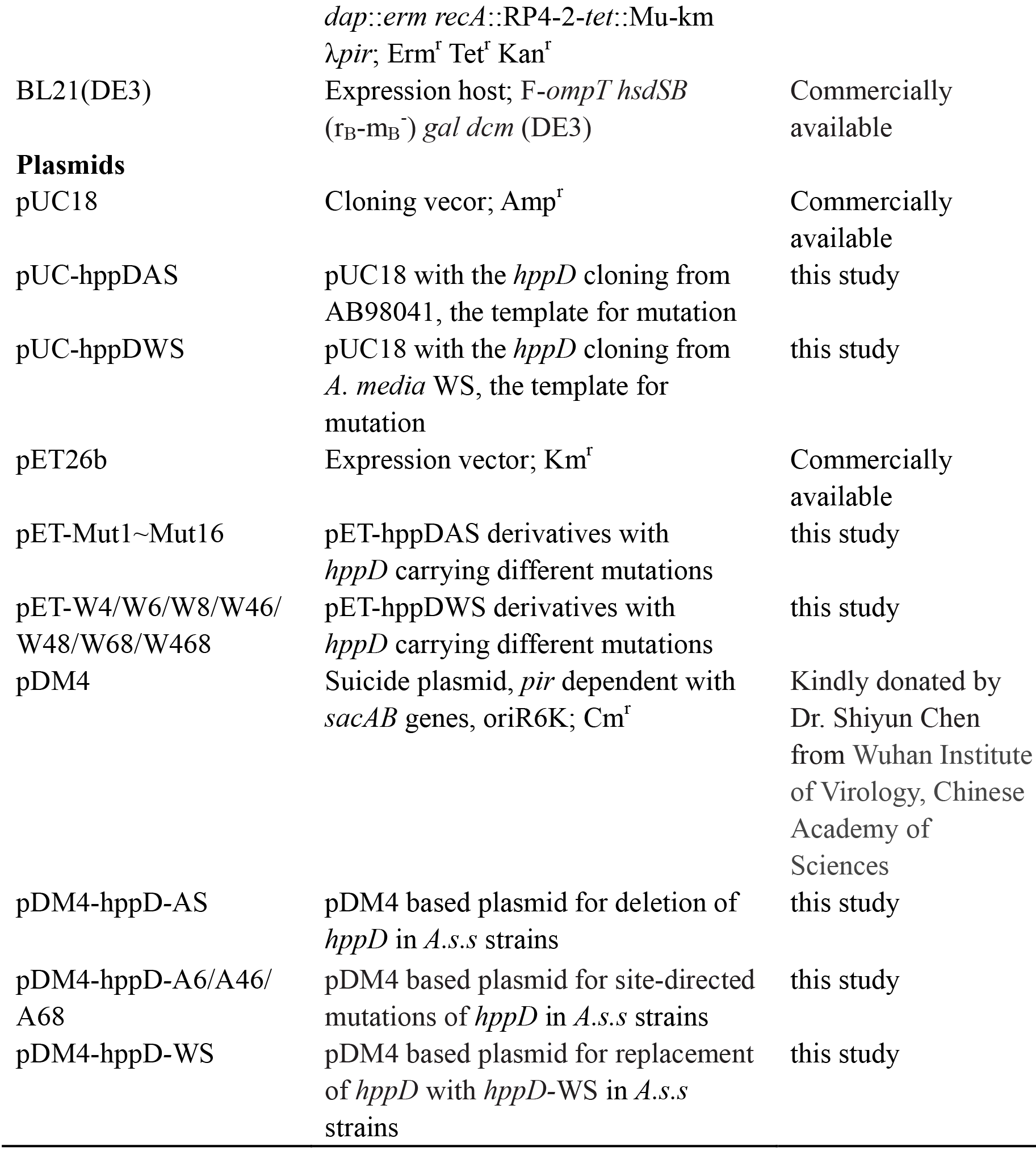

### Identification of residues in HppD-AS for site-directed mutagenesis

HppD is the most important enzyme in the HGA-melanin synthesis pathway. Almost all *Aeromonas* species for which sequence data has been submitted to GenBank have homologous *hppD* genes, regardless of pigment production. Since thermosensitive pigmentation does not occur in other *Aeromonas* species and the expression of HppD alone is sufficient for this phenotype, we hypothesized that comparing HppD-AS with HppD from other *Aeromonas* species would reveal the crucial residues that are important for HppD-AS thermolability. We used the “consensus approach” to identify potential amino acids for mutagenesis. This approach is based on replacing a specific amino acid with the most frequent amino acid at a given position based on the alignment of homologous proteins, which has been applied successfully for many protein engineering studies (18–20). Fifteen homologous HppDs from different representative *Aeromonas* species were selected for sequence alignments. These all shared greater than 80% amino acid identities with that of HppD-AS. As shown in Fig. 1, sequence analysis revealed a total of eleven positions in HppD-AS that differed from the most conserved amino acids in homologous proteins. These residues were found to be S5, S10, S18, A96, P103, T107, L119, T160, S298, D301, and Q303. In addition, we found five other residues that are present in nearly half of the homologues, specifically S8, Q180, S232, L255, and Q262 (Fig. 1). Therefore, we replaced all sixteen of these residues in HppD-AS with the most prevalent amino acid at that position. Subsequently, single variants (S5T, S8A, S10T, S18T, A96S, P103Q, T107A, L119P, T160A, Q180H, S232A, L255M, Q262H, S298A, D301E, and Q303R) were generated and termed Mut1 to Mut16, respectively.

**Figure 1.**
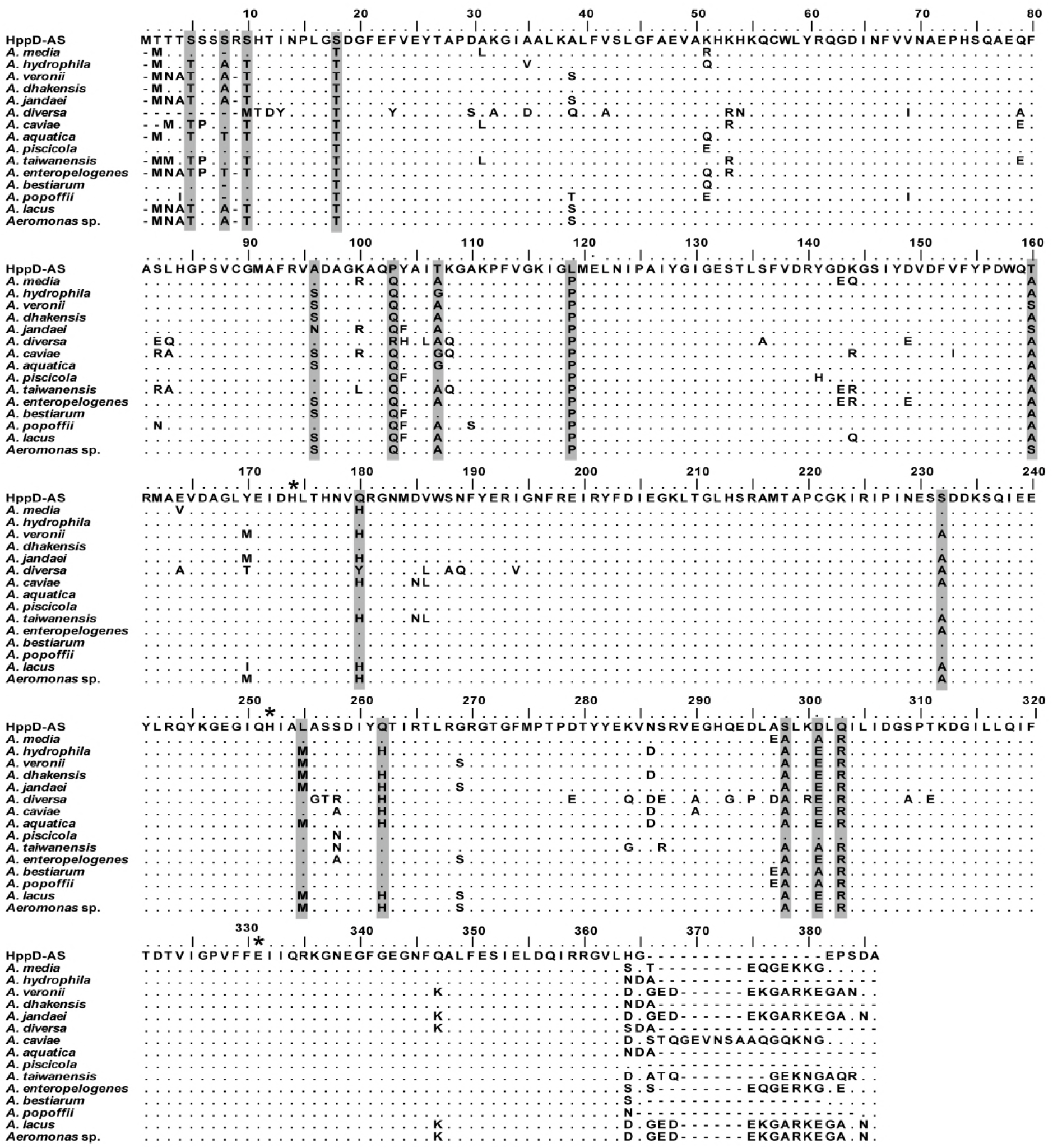
Amino acid sequence alignment of HppD from *A.s.s* and other fifteen *Aeromonas* species. The sequences were aligned using the ClustalW algorithm. The identical residues are plotted as dots. The mutant positions selected are highlighted in gray. Asterisks mark the catalytic triad residues. The numbers above the sequences refer to the amino acid positions in HppD-AS. The full strain names and accession numbers of HppD are listed in Table S2.

### Identification of residues in HppD-AS that are critical for temperature-dependent pigmentation in recombinant *E. coli* strains

As heterologous expression of *hppD*-AS in *E. coli* BL21 results in pigmentation at 22 °C but not at ≥ 30 °C, we tested whether any of our HppD-AS mutants resulted in melanin production at ≥ 30 °C in recombinant *E. coli* strains. *hppD*-WS (*hppD* from *A. media* strain WS) and empty plasmid were used as controls. As presented in Fig. 2A, compared to that in the strain harboring an empty plasmid, all the recombinant *E. coli* strains expressing HppD could synthesize pigment at 22 °C, indicating that all *hppD* variants were functional. As expected, BL21(pET26-*hppD*-AS) lost melanogenic ability when the induction temperature was increased to 30 °C or 37 °C, whereas BL21(pET26-*hppD*-WS), which expressed HppD-WS, still formed pigment. Interestingly, when analyzing the sixteen HppD-AS mutants, *E. coli* expressing three variants (Mut4 [S18T], Mut6 [P103Q], and Mut8 [L119P]) produced pigment at 30 °C (Fig. 2A, which only shows partial results of mutants), and Mut6 was even found to result in pigment accumulation at 37 °C.

To further confirm the role of these three residues in the thermosensitive melanogenesis phenomenon, double mutants Mut46 (S18T/P103Q), Mut48 (S18T/L119P), and Mut68 (P103Q/L119P), as well as a triple mutant Mut468 (S18T/P103Q/L119P) were generated and studied. As shown in Fig. 2B, any combination of these three mutations enabled recombinant *E. coli* strains to form melanin at 30 °C. Notably, any mutant containing the P103Q substitution (Mut46, Mut68, and Mut468) could generate a significant amount of pigment at 37 °C, comparable to that with *hppD*-WS. The double mutant Mut48 also resulted in a slight sign of melanization at 37 °C, whereas the single mutant Mut4 or Mut8 did not. These data demonstrate that the effect of each mutation is additive. Taken together, these results indicate that the three substitutions (S18T, P103Q, and L119P) in HppD-AS significantly increase heat resistance of melanogenesis in recombinant expression strains, of which Mut6 (P103Q) has the strongest effect.

**Figure 2.**
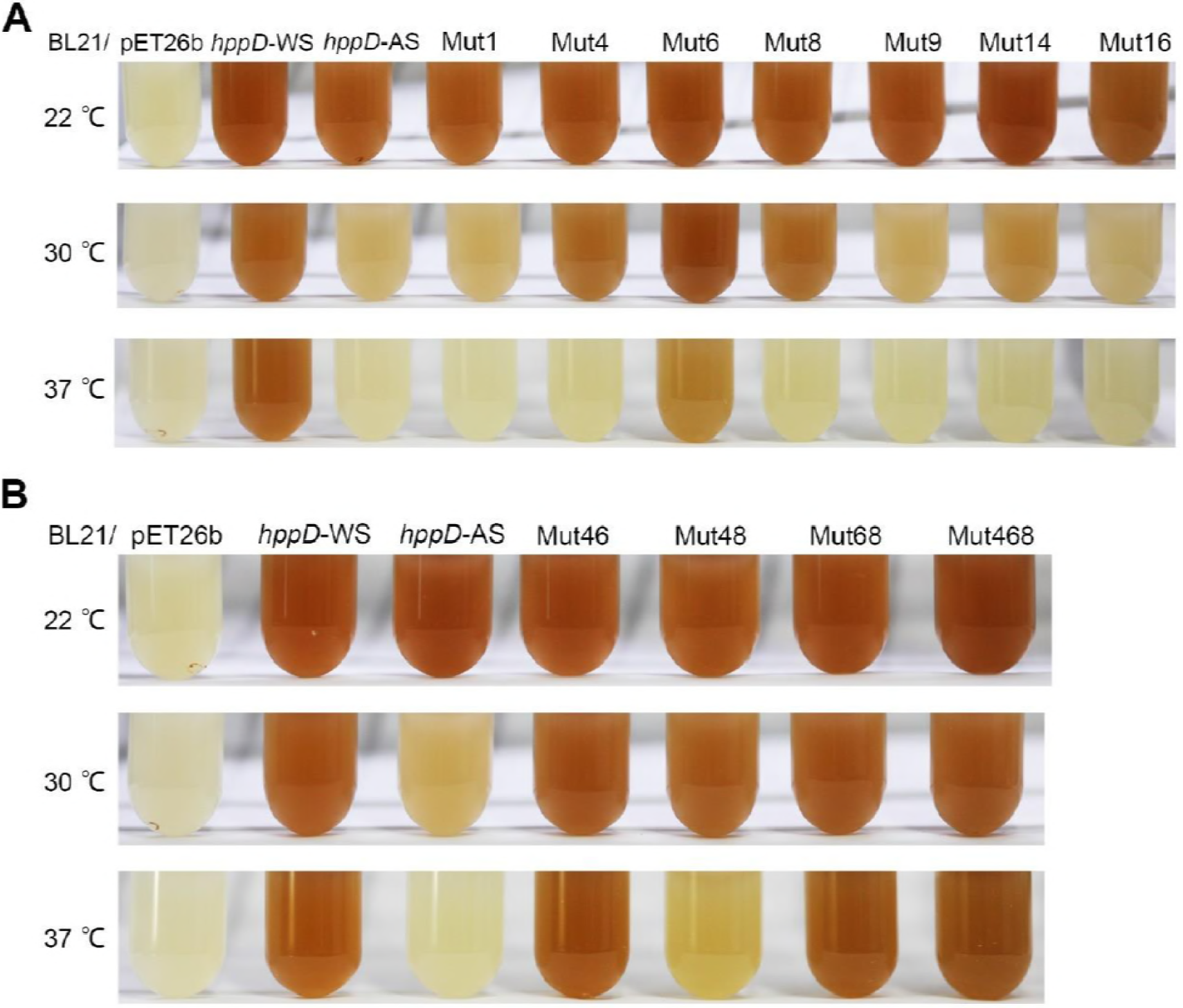
The melanin production of *E. coli* strains expressing *hppD*-AS variants. Recombinant *E. coli* strains expressing the single mutants (**A**) and double/triple mutants (**B**) were cultured at 22, 30, and 37 °C after the induction of IPTG. *E. coli* strains containing the empty vector or expressing wild type *hppD*-AS and *hppD*-WS were served as controls.

The aforementioned results clearly demonstrate a key role for three residues in HppD-AS in temperature-dependent pigmentation. We suggested that in theory, replacing the corresponding residues of HppD from other *Aeromonas* species that undergo temperature-independent pigmentation with those of HppD-AS should result in temperature-sensitive melanization. We thus substituted the corresponding residues of HppD-WS with the amino acids of HppD-AS at positions S18, P103, and L119. As a result, single point mutants W4(T16S), W6(Q101P) W8(P117L), and double/triple mutants W46(T16S/Q101P), W48(T16S/P117L), W68 (Q101P/P117L), and W468(T16S/Q101P/P117L) of HppD-WS were constructed and tested. As shown in Fig. 3, cells expressing the W4 or W8 mutant formed pigment at all tested temperatures, suggesting that these mutations do not dramatically affect the function of HppD-WS. However, cells expressing the W6 mutant failed to produce pigment at 37 °C, consistent with its important role in temperature-dependent pigmentation. Similar to the observed additive effect of these mutations (mentioned previously herein), strains expressing any of the double mutants failed to synthesize pigment at 37 °C. Importantly, cells expressing the triple mutant W468 produced less pigment at 22 °C, compared to that in other strains, and completely lost melanin formation ability at 30 °C and 37 °C. This indicated that we had constructed an HppD-AS-like HppD-WS protein by altering only three residues.

**Figure 3.**
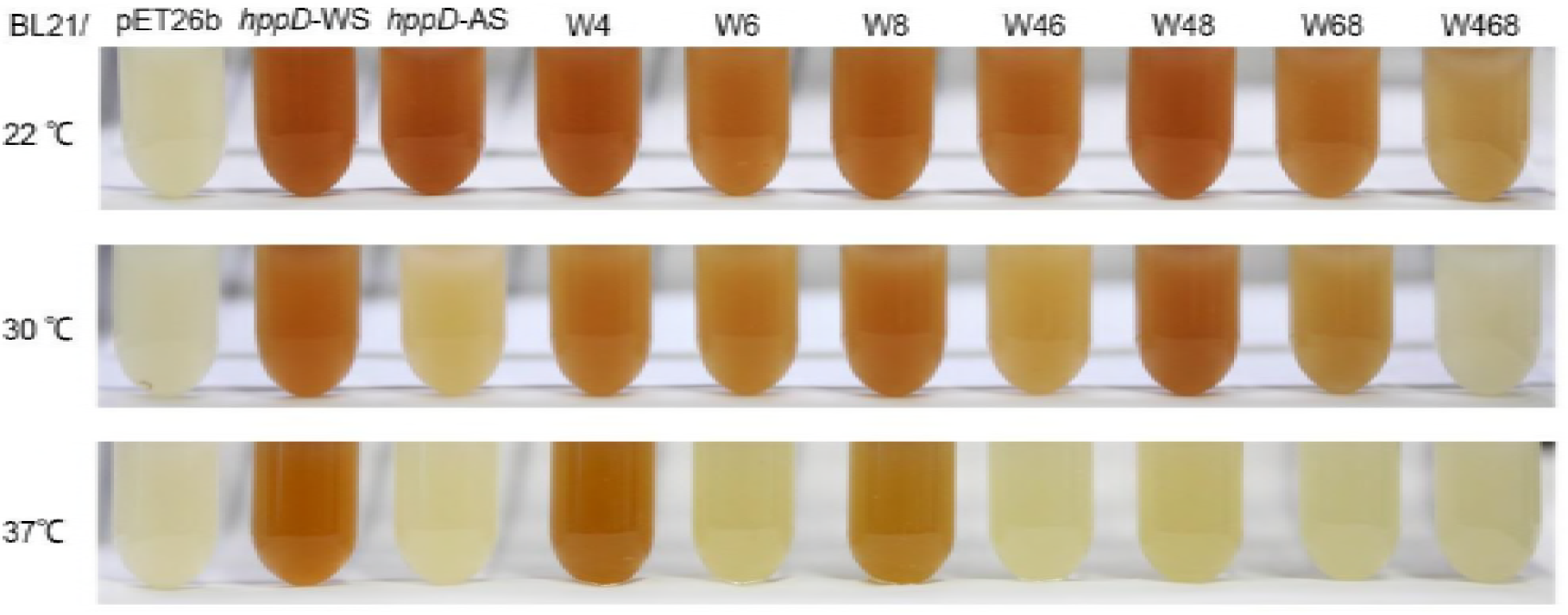
The melanin synthesis phenotype of *E. coli* strains expressing *hppD*-WS variants. The corresponding residues of HppD-WS were replaced with the amino acids of HppD-AS. Strains were induced by IPTG at different temperatures (22, 30, and 37 °C, respectively). *E. coli* strains carrying an empty vector or expressing wild type *hppD*-AS and *hppD*-WS were employed as controls.

### Verification of the critical role of HppD in temperature-dependent pigmentation in *A.s.s*

Using the heterogeneous *E. coli* system, we established that temperature-dependent melanogenesis in *A.s.s* is due to the function of HppD-AS at different temperatures and identified critical residues of HppD-AS that are involved in this phenotype. However, drawing conclusions from this artificial system is dangerous unless we could prove it in the natural biological system. To verify that our findings in *E. coli* are relevant for *A.s.s*, we modified chromosomal *hppD* in *A.s.s* strains. We first knocked out the *hppD* gene in wild-type *A.s.s* strains to determine if this would abolish pigmentation. As expected, the three *hppD* deletion mutants (AB98041Δ*hppD*, 2013-8Δ*hppD*, and 2014-235Δ*hppD*) all completely lost their ability to generate pigment regardless of the culture temperature (Fig. 4). This confirmed that pigmented *A.s.s* strains mainly synthesize pyomelanin through the HGA pathway. We next constructed AB98041 mutant strains carrying the *hppD* mutant alleles that eliminated temperature-dependent pigmentation in *E. coli* using the AB98041Δ*hppD* strain as the background. Three mutant strains, AB98041(*hppD*-6 [P103Q]), AB98041(*hppD*-46 [S18T/P103Q]), and AB98041(*hppD*-68 [P103Q/L119P]), were generated and tested for pigmentation at different temperatures. As shown in Fig. 4A, unlike the wild-type strain, AB98041(*hppD*-6), AB98041(*hppD*-46), and AB98041(*hppD*-68) were able to produce pigment at 30 °C. Furthermore, when the *hppD* gene of AB98041 was replaced with hppD-WS, the resultant strain, designated AB98041(*hppD*-WS), synthesized melanin at 30 °C (Fig. 4A). Similar phenotypes were observed with two other *A.s.s* isolates when their *hppD* genes were replaced with *hppD*-WS, respectively (2013-8[*hppD*-WS] and 2014-235[*hppD*-WS]) (Fig. 4B and 4C). Based on these results, we ascertained that the inhibition of pigmentation in *A.s.s* strains at elevated temperatures is caused by the diminished function of HppD.

**Figure 4.**
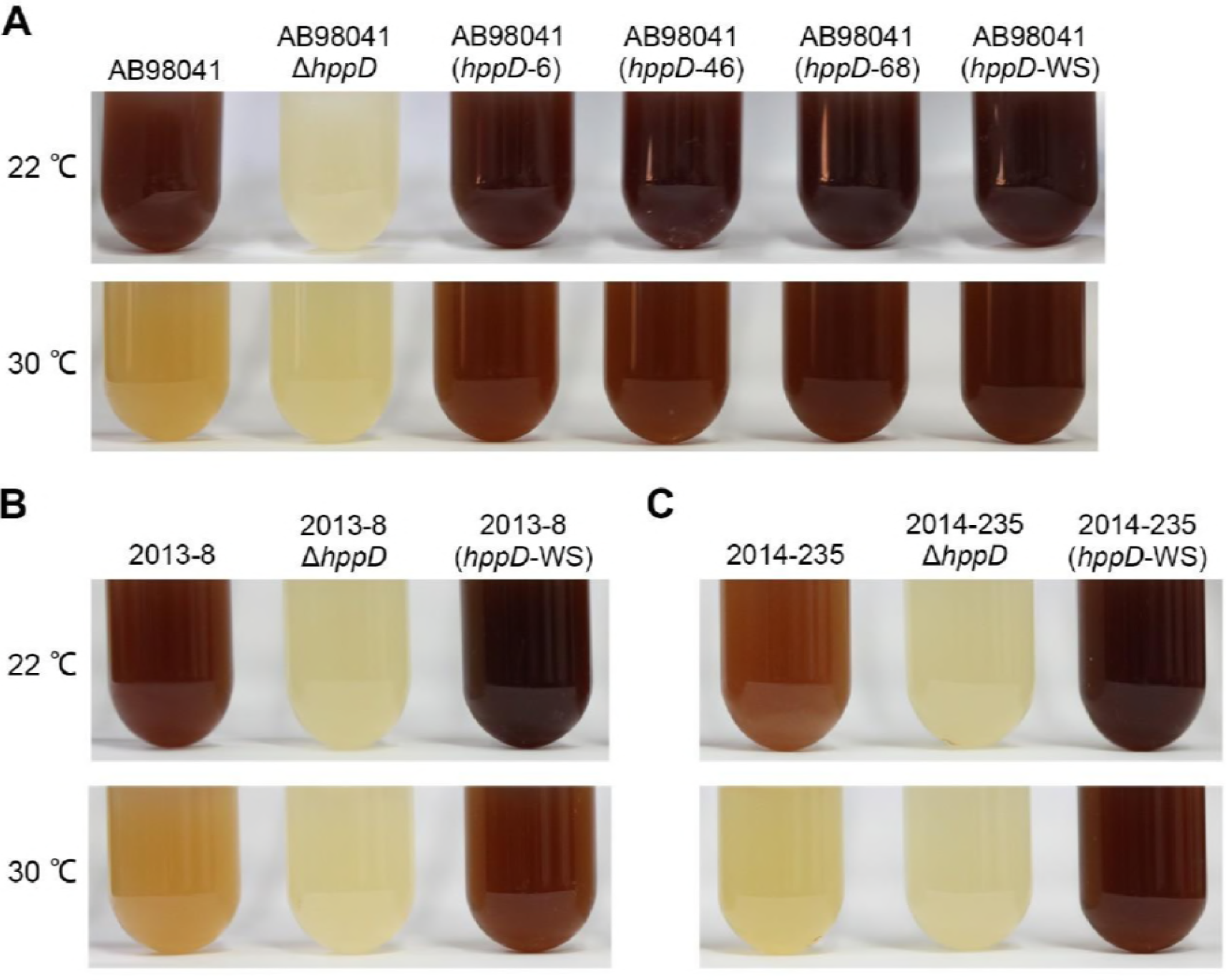
*hppD*-AS mutants result in temperature-independent melanogenesis in *A.s.s* strains. Shown are pigmentation phenotypes of wild-type strain AB98041 and its *hppD* mutants **(A)**, wild-type strain 2013-8 and its *hppD* mutants **(B)**, wild-type strain 2014-235 and its *hppD* mutants **(C)**, which were cultivated in LB medium for 48 h at 22 and 30 °C, respectively.

### HppD-AS is thermolabile *in vitro*

The previously mentioned results suggest that thermosensitive melanization in *A.s.s* is likely due to the thermolability of HppD-AS, and that mutations that stabilize HppD-AS result in a temperature-independent phenotype. To confirm this, wild-type HppD-AS, HppD-WS, and seven HppD-AS variants (Mut4, Mut6, Mut8, Mut46, Mut48, Mut68, and Mut468) that resulted in the melanization of *E. coli* at 30 °C were homogenously purified and characterized as described in the Materials and Methods. Based on sodium dodecyl sulfate-polyacrylamide gel electrophoresis (SDS-PAGE), all purified mutant enzymes appeared as a single band with the same molecular mass (~42 kDa) as HppD-AS, in agreement with the predicted molecular mass of 41227.47 Da (Fig. S2).

The effect of temperature on the activities of wild-type and the seven HppD-AS mutant enzymes was determined at temperatures ranging from 16 to 53 °C. To our surprise, purified HppD-AS displayed no activity at any temperature. Moreover, we found that the HppD-AS crude extract lost enzyme activity at 4 °C within 8-10 h probably due to its excessive thermolability. Therefore, the enzymatic activity and thermal stability of HppD-AS was assayed using its fresh crude extract, whereas purified enzymes were used for all other mutants. As shown in Fig. 5A and Fig. S3, HppD-AS showed maximum activity at 22 °C. However, mutant enzymes such as Mut6 and Mut468 exhibited higher optimum temperatures of 30 °C and 37 °C, respectively. HppD-WS displayed the highest activity at 37 °C. And at 45 °C, wild-type HppD-AS exhibited only 43% of its maximal activity, whereas Mut8, Mut6, and Mut468 mutants retained around 51%, 94%, and 92% of their individual maximum activities, respectively. To evaluate the thermostability of each enzyme, protein was incubated at 40 °C and after various incubation times, samples were taken to determine residual activity. As shown in Fig. 5B, HppD-AS activity was highly sensitive to thermal inactivation. The residual activity of HppD-AS decreased exponentially to only 8% of initial activity within 10 min, whereas the activity of HppD-WS was not significantly reduced even after incubation for 80 min. However, Mut4, Mut8, and the double mutant Mut48 exhibited longer half-lives than wild-type HppD-AS, indicating that these two mutations contributed to enhancing the thermostability of HppD-AS. Remarkably, the substitution P103Q obviously improved protein stability. The single mutant Mut6 retained approximately 75% of initial activity after incubation at 40 °C for 80 min, and combining this variant with the other two mutations further increased enzyme stability, as indicated by the activity of the triple mutant Mut468. These results together demonstrate that wild-type HppD-AS is extremely thermolabile and that substitutions at three residues (S18T, P103Q, L119P) can substantially increase enzyme thermostability such that *A.s.s* strains containing these mutated *hppD* genes can produce melanin at 30 °C. In other words, HppD thermolability is suggested to mediate the abolished pyomelanin formation of *A.s.s* strains at higher temperatures.

**Figure 5.**
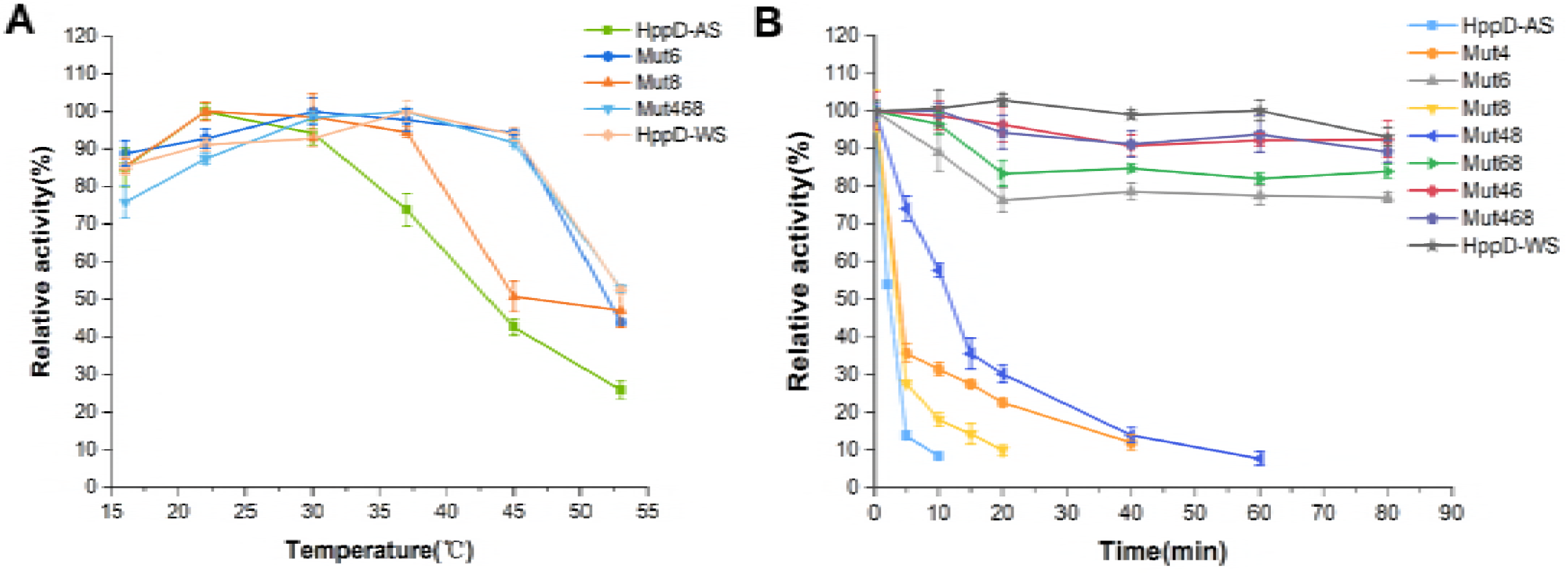
Effect of temperature on the activities and stabilities of wild-type and mutant HppD enzymes. **(A)** Optimum temperatures of several representative enzymes. Activity was measured in 100 mM Tris-HCl (pH 7.5) at 16, 22, 30, 37, 45, and 53 °C for 30 min, respectively. The total activity at the optimum temperature is set as 100%. **(B)** Thermostabilities of HppD enzymes. The proteins were incubated at 40 °C, and aliquots were then taken at different time points to determine residual activities at 22 °C. The activity of each enzyme is normalized to its activity prior to incubation, which is defined as 100%. The error bars show the standard deviations of triplicate assays.

### Structural analysis of HppD-AS mutation sites

To gain insight into the mechanisms of enhanced thermostability mediated by the three mutations (S18T, P103Q, L119P) in HppD-AS, we constructed a structural model of wild-type HppD-AS based on the crystal structure of HppD from *Pseudomonas fluorescens* (PDB ID, 1CJX) (Fig. 6). For this, the three substituted residues were mapped to the generated model structure. All three residues were found to be located in the N-terminal domain of the enzyme and the effect of these substitutions on HppD thermostability was then evaluated (Fig. 6).

**Figure 6.**
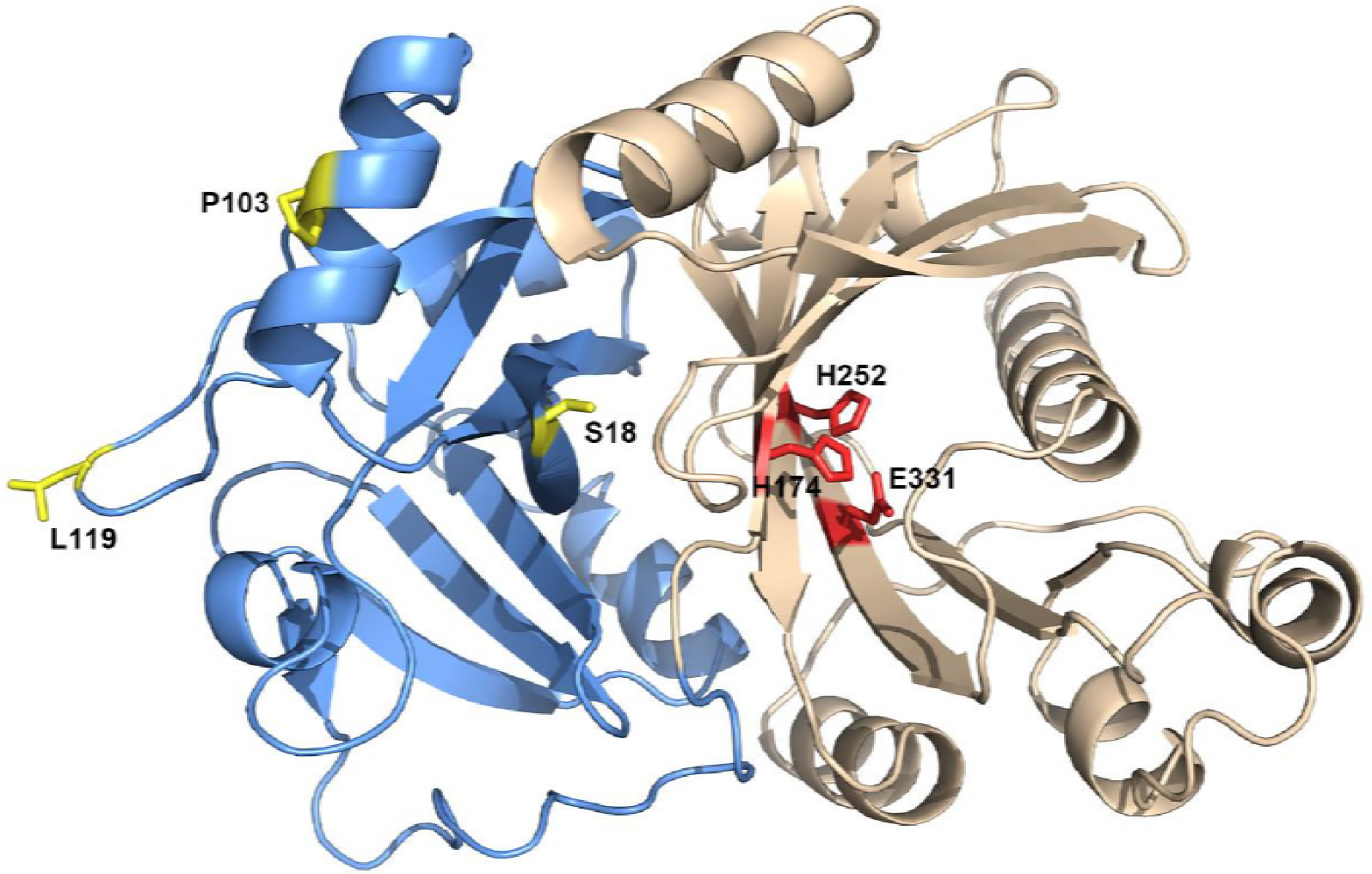
The structural model of wild-type HppD-AS. The structural model of wild-type HppD-AS. The model structure was generated based on the HppD structure of *Pseudomonas fluorescens* (PDB ID, 1CJX). The two barrel-like domains, N-terminal domain and C-terminal domain, are depicted in blue and beige, respectively. The catalytic triad residues (H174, H252, and E331) are highlighted by red sticks. The three substituted residues (S18, P103, and L119) are marked in yellow.

The S18 residue resides in the first β strand (β1) of the enzyme, which forms a β-sheet with the parallel β4 strand and antiparallel β5 strand (Fig. 7A). The S18T substitution was determined to be a very conservative change because both Ser and Thr are uncharged polar amino acids that differ only by one methyl group. It was predicted that this mutation would not cause significant changes to the local structure or the number of hydrogen bonds (Fig. 7B). However, it has been reported that Thr has a greater intrinsic propensity to form β-sheets than Ser, which was verified by both statistical analysis of proteins with known structures (21, 22) and experimental measurement using different host-guest model systems (23). The distinct propensities to form β-sheets were considered due to different mean curvature parameters of each amino acid (24). As Thr has a larger accessible area in its side chain relative to that of Ser, which results in a smaller mean curvature value in the backbone surface, it can readily adopt a β-sheet pattern and stabilize β-sheets (24). Thus, the enhanced stability of the S18T mutant of HppD-AS is likely due to a slightly more stable structure. Nonetheless, the effect of the mutation was predicted to be moderate, consistent with the moderate increase in pigmentation at higher temperatures conferred by the mutation *in vivo*.

The substitution P103Q resulted in temperature-independent pigmentation in *A.s.s in vivo* and had the greatest effect on enzyme stability *in vitro*. Based on the modeled structure, P103 is located in the internal α3 helix (Fig. 7C). It is well known that Pro is a strong breaker of α helical structure and has a low tendency to form α-helices (21, 25, 26). Due to the pyrrolidine ring in its side chain, Pro has no amide hydrogen to form hydrogen bonds while it is in the middle of a polypeptide chain. Thus, Pro103 cannot participate in hydrogen bonding to the carbonyl oxygen atom of the preceding fourth residue Gly99 in the α-helix (Fig. 7C). Furthermore, it has been widely proven that the presence of a proline residue in α-helices can cause a pronounced kink between the segments preceding and following proline (27, 28). This kink results from the reduced number of helix backbone hydrogen bonds and helps to avoid steric clash between the pyrrolidine ring and the local backbone (28). As a consequence, the presence of proline would greatly increase the flexibility of helices. However, substituting Pro103 with Gln would eliminate the distortion in the helical structure caused by the pyrrolidine ring of proline. Furthermore, glutamine contains two amide protons that would introduce two new hydrogen bonds with the carbonyl oxygen of Gly99 (Fig. 7D). These changes are suggested to significantly increase the rigidity of the α3 helix, thus stabilizing the overall structure of the protein. This is consistent with the potent effect of this mutation on the activity of HppD-AS *in vivo* and *in vitro*.

L119 was found to be situated at the second residue of a β-turn between strands β6 and β7 (Fig. 7E). The reason for increased thermostability of the L119P mutant enzyme appears to lie in a reduction in the flexibility of the β-turn local backbone. As proline adopts fewer conformations in its backbone than other amino acids because of the rigid pyrrolidine ring, it can constrain the conformational freedom of Cα-N rotation (29) and stabilize the protein by reducing the entropy of the denatured state (30). The introduction of a proline into flexible regions to improve the thermostability of proteins has been well documented (31–33). Certain key positions for protein thermostabilization by proline have also been declared, namely, flexible loops, the N-caps of α-helices, and the first or second site of β-turns (33). In addition, except for the hydrogen bonds formed by its oxygen atom with K53, L119 is not involved in other H-bonds via its backbone amide (Fig. 7E). Therefore, the L119P mutation would make the β-turn more rigid without negative side effects. According to the above analysis, we concluded that all three substitutions (S18T, P103Q, and L119P) stabilize the structure of the protein by strengthening secondary structural elements of HppD-AS.

**Figure 7.**
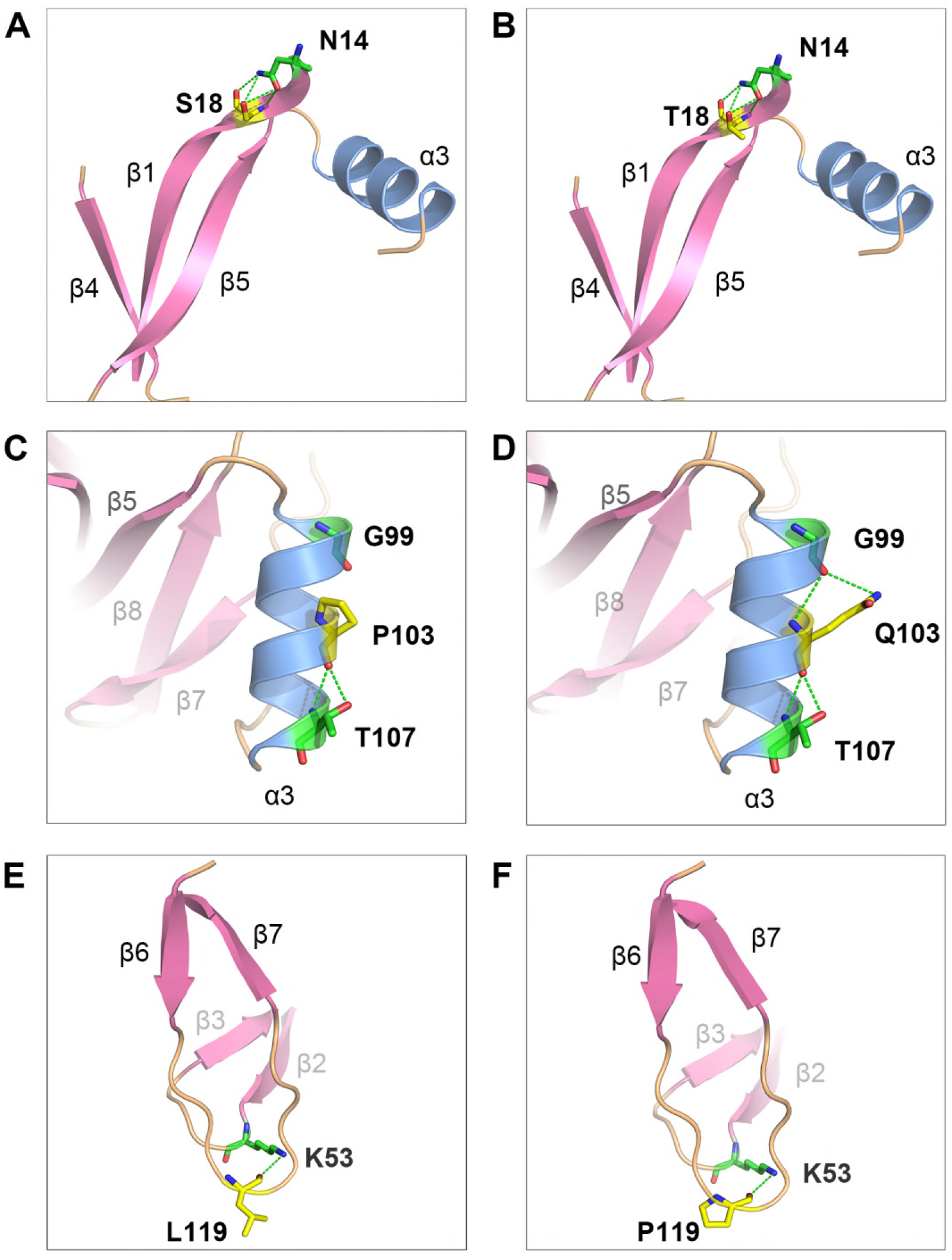
Potential impact of the substitutions on the structure of HppD-AS. Potential impact of the substitutions on the structure of HppD-AS. Shown are close-up views of local structures around the positions 18, 103, and 119 before (**A**, **C**, **E**) and after (**B**, **D**, **F**) mutations, respectively. The original and substituted residues are indicated by yellow sticks. The green dashed lines represent hydrogen bonds.

## Discussion

The temperature-sensitive melanization of *A.s.s* strains has been observed for many years. However, since very little is understood about the associated melanin synthesis pathway, the mechanism underlying this phenomenon remains unclear. We previously showed that *A. media* WS produces pyomelanin through the HGA pathway. Moreover, we found that heterologous expression of HppD from an *A.s.s* strain in *E. coli* resulted in temperature-sensitive pigmentation, indicating that *A.s.s* strains likely produce melanin through the HGA pathway and that HppD plays a critical role in the thermal sensitive melanization process. Here, we confirmed that HppD is critical for melanization in *A.s.s* and demonstrated that the extreme thermolability of HppD-AS is responsible for the temperature-dependent melanogenesis. Here, we also identified the critical residues of HppD-AS that are important for its thermolability and showed that replacing the corresponding residues in HppD from *A. media* WS results in thermolabile enzymes.

Among the three identified residues, S18 and L119 were shown to have a moderate effect, whereas P103 had the strongest impact on thermal lability. A single P103Q substitution greatly increased the thermostability of HppD-AS and was sufficient to eliminate the thermo-sensitive characteristic in both recombinant *E. coli* and *A.s.s* strains. This indicates that mutation in only one site can substantially effect protein conformation, thus changing the property of the protein dramatically. Through homology modeling, we revealed that this site resides in the sixth position of an α-helix in HppD-AS. The proline residue is rarely found after the fourth residue in α-helices of globular proteins because of its ability to disrupt α-helical structure (25). This should be the main reason for the thermolability of HppD-AS. However, why would such a seemingly unfavorable residue has been reserved in HppD-AS?

We aligned the HppD-AS sequence with its homologs from all other *A. salmonicida* subspecies and *Aeromonas* species published in GenBank to examine the degree of conservation of this site. Intraspecific sequence alignment was conducted among isolates including *A. salmonicida* subsp. *masoucida* RFAS1, subsp. *pectinolytica* 34mel^T^, and nine un-subspeciated strains (see Table S1 for detailed information). Alignment results showed that there are some residue variations in the HppD of other *A. salmonicida* subspecies compared to the sequence of HppD-AS (Fig. S4). Notably, among the variant positions, Pro103 and Leu119 of HppD-AS were found to be absolutely conserved and clearly distinct from the Gln and Pro in other subspecies, respectively. More strikingly, interspecific sequence comparison with totally 206 recognized and published *Aeromonas* spp. isolates found that Pro and Leu at these two sites were only conserved within *A.s.s* isolates and that in other bacterial species, these were substituted with different residues including Gln, Leu, Arg, and Pro (data not shown). As *A.s.s* are strict psychrophilic bacteria, we surmise that the consistent preservation of Pro103 and Leu119 in HppD from all *A.s.s* strains is a protein adaptation that occurred during evolution to adopt a psychrophilic lifestyle. The wide distribution of such substitutions might imply their advantage. The assessment of the activities of each enzyme at various temperatures showed that the optimal temperature of HppD-AS was 22 °C, whereas that of other mutants gradually shifted from 22 to 37 °C, and HppD-WS had maximum activity at 37 °C (Fig. 5A). This agrees with a previous study suggesting that psychrophilic enzymes usually possess high catalytic activity at the optimal growth temperatures of bacterium (34). It indicates that although Pro113 and Leu119 residues decrease thermostability, they substantially improve the catalytic activity of HppD-AS at low temperatures. This is in accordance with the common strategy of cold adaptation for proteins; specifically, psychrophilic enzymes always have amino acid substitution preferences that avoid forming helical structures and participating in local interactions to confer higher protein flexibility to offset the slow catalytic reaction rate inherent to low temperatures (34, 35). Furthermore, the high conservation of *hppD* sequences among *A.s.s* strains might be attributed to the clonal population structure of this subspecies (36). Such high clonality is probably an adaptation to the particular host, salmonid fish, in which clonal outbreaks are prone to occur (37).

Although *A. salmonicida* species are classified into five subspecies including *salmonicida*, *achromogenes*, *smithia*, *masoucida*, and *pectinolytica*, as reported in Bergey’s Manual of Systematic Bacteriology (10), many laboratories currently designate subspecies *salmonicida* as “typical” and any isolate diverging phenotypically and molecularly as “atypical” strains. The typical isolates comprise an extremely compact group and are considered clonal (38), whereas atypical isolates are more heterogeneous and show a wide range of hosts including non-salmonid fish as well as salmonids (39). The convoluted history of taxonomy and nomenclature for *A. salmonicida* species makes the clear assignment of strains difficult. Concerning the particular possession of Pro103 and Leu119 in HppD-AS, these residues could be used as potential molecular markers for identifying *A.s.s* isolates. The conventional and readily observable feature to differentiate subspecies *salmonicida* from atypical strains is the production of a brown diffusible pigment. However, this rule was not always applicable; for example, some *A.s.s* isolates lack pigment production and some atypical isolates synthesize melanin (40–42). Other biochemical tests such as indole production and gluconate oxidation have similar caveats (43). A variety of molecular genetic techniques have also been applied for the characterization of typical and atypical *A. salmonicida* strains, including DNA:DNA reassociation (44), ribotyping (45), PCR-based typing analysis (46), plasmid profiling (47), pulse-field gel electrophoresis (48), amplified fragment length polymorphism analysis (49), and IS-based RFLP (50), among others. Nonetheless, these methods are often time-consuming and occasionally controversial. DNA sequencing analysis based on the 16S rRNA gene also makes the identification challenging at the species level because of the extreme intraspecies conservation of this gene (51). Recently, *vapA*, a gene encoding the A-layer, was used to establish sequence schemes and to distinguish typical *A. salmonicida* from atypical isolates, with partial nucleotide sequence homogeneity between the former but heterogeneity among the latter (52). However, it has been shown that there can be a high frequency of mutations in *vapA* when the strains are cultured in unfavorable conditions. The identical *hppD* nucleotide sequence described here, and the distinct presence of Pro103 and Leu119 residues compared to its homologs from other *Aeromonas* species, provide a new pragmatic parameter to easily discriminate *A.s.s* from atypical strains and other *Aeromonas* species. Furthermore, this could complement the A-layer typing method, which cannot classify strains lacking the A-layer (e.g. subspecies *pectinolytica*) (52).

It has been generally reported that inactivation of homogentisate 1,2-dioxygenase (HmgA), an enzyme that degrades HGA into Kreb’s cycle, can result in the accumulation of pyomelanin (53–55). By examining the *hmgA* genes from laboratorial and published *A.s.s* strains, we observed that they all have frameshift mutations and form pseudogenes. This suggests that all of these *A.s.s* isolates are pyomelanin-producing. Moreover, the presence of the conserved Pro103 residue in HppD protein implies that the temperature susceptibility of melanization is exclusive and ubiquitous for *A.s.s.* As reported previously, the genome of *A.s.s* A449 revealed a significant number of pseudogenes, including many virulence genes, genes encoding transcription factors, and genes encoding basic metabolic enzymes, among others (56). Since *A.s.s* is a specific pathogen of *Salmonidae*, the accumulation of pseudogenes has been suggested to be a hallmark of adaptation to the specific host (56). Considering that melanin has been associated with virulence in many pathogenic microorganisms, it is tempting to speculate that both disruption of the *hmgA* genes and temperature-sensitive pigmentation represent pathoadaptation towards particular salmonid hosts. However, the correlation between melanin and host specificity has not been reported. Subsequently, further pathogenicity experiments are required to validate this contention.

## Materials and Methods

### Bacterial strains, plasmids, and growth conditions

All the parental and derivative strains and plasmids used in this study are listed in Table 1. *A.s.s* strain AB98041 was obtained from the China Center for Type Culture Collection (CCTCC) and *A.s.s* strain KACC14791 was purchased from the Korean Agricultural Culture Collection (KACC). The other two *A.s.s* isolates, 2013-8 and 2014-235, were kindly donated by Dr. Tongyan Lu from Heilongjiang River Fisheries Research Institute, Chinese Academy of Fishery Sciences. *E. coli* strains were routinely cultured in Luria-Bertani (LB) broth or agar at 37 °C unless otherwise indicated. Heterologous expression strains were also grown at 22 or 30 °C to compare melanin production. The wild-type and mutant *A.s.s* strains were grown in LB medium at 22 or 30 °C to observe the melanin synthesis phenotype. Antibiotics were supplemented in the medium at the following concentrations when necessary: ampicillin (100 μg/ml), kanamycin (50 μg/ml), chloramphenicol (34 μg/ml). For cultivation of *E. coli* strain β2155, 100 μg/ml diaminopimelic acid (DAP) was added.

### Site-directed mutagenesis and heterologous expression of the *hppD* gene

The *hppD* genes from *A.s.s* AB98041 and *A. media* WS were previously cloned using the primer pairs hppD(AS)-S/hppD(AS)-A and hppD(WS)-S/hppD(WS)-A (7). The amplified fragments were then digested with *Nde*I and *Hind*III (Thermo Fisher Scientific, USA) and inserted into the pUC18 vector (TaKaRa, Japan) digested with the same enzymes to construct plasmids pUC-hppDAS and pUC-hppDWS, respectively. The double-stranded, dam-methylated recombinant plasmids were isolated from *E. coli* DH5α, confirmed by DNA sequencing, and used as the PCR template to construct all mutants. Mutagenesis was extended by KOD-Plus-Neo DNA polymerase (TOYOBO, Japan) using a pair of mutagenic primers containing the appropriate base changes (Table S3). The products were treated with *Dpn*I (Thermo Fisher Scientific, USA) at 37 °C for 30 min, and nicked plasmid DNA was transformed into *E. coli* DH5α for automatic repair. The plasmids harboring expected mutations were identified by restriction analysis and DNA sequencing. The mutated fragments were digested and ligated into expression vector pET26b (Novagen, Germany) at *Nde*I and *Hind*III restriction sites. The ligation products were transformed into *E. coli* BL21 (DE3), which were selected for transformants. Correct transformants, confirmed by PCR, were cultivated at 37 °C overnight in 5 ml LB medium containing 50 μg/ml kanamycin, and then transferred to the same fresh medium for further culture until the optical density at 600 nm (OD_600_) reached ~0.6. Isopropyl β-D-thiogalactopyranoside (IPTG) was added to a final concentration of 0.5 mM to induce enzyme expression. Subsequently, cultures were incubated at the indicated temperatures (22, 30, and 37 °C, respectively) for another 36 h to examine the production of melanin.

### Construction of *A.s.s* strains with different *hppD* alleles

Deletion, site-directed mutation, and replacement of the *hppD* gene in *A.s.s* AB98041 were all performed by the double homologous recombination method. To construct the AB98041 *hppD* deletion strain, DNA fragments carrying 500 bp regions upstream and downstream of the *hppD* open reading frame were amplified from the chromosomal DNA of AB98041 with the primer pairs hppD(up)-F/hppD(up)-R and hppD(dn)-F/hppD(dn)-R (Table S3), respectively. Upstream and downstream fragments were spliced by overlap-extension PCR via the overlapping homologous sequences of hppD(up)-R and hppD(dn)-F. The assembled fragment was digested with *Xba*I and *Sma*I and cloned into the suicide vector pDM4 containing the *sacB* gene. The resultant plasmid pDM4-hppDAS was transformed into the *E. coli* β2155(*λ*pir) strain, which can grow only in the presence of DAP (57). Two-parental mating was performed to transfer the recombinant plasmid pDM4-hppDAS into AB98041. Briefly, the AB98041 recipient strain and the β2155 (harboring pDM4-hppDAS) donor strain were grown overnight in LB medium and LB-chloramphenicol medium containing 100 μg/ml DAP, respectively. Cultures were mixed in a 200 μl volume at a ratio of 1:1, centrifuged at 8,000 × *g* for 5 min, washed twice with 400 μl of 10% glycerol, resuspended in 50 μl of 10% glycerol, and then spotted on LB-DAP agar plates. After incubation at 22 °C for 12 h, cells were harvested and resuspended in 2 ml of LB. Fifty microliters of culture was then plated on LB-chloramphenicol agar to select single recombinants. Colonies were transferred to LB agar plates and grown for 24 h for a second allelic exchange, and then transferred to LB agar plates containing 15% sucrose. Double recombinants that could grow in the presence of sucrose were identified. Potential *hppD* deletion mutants were confirmed by PCR analysis and DNA sequencing.

To construct *hppD* mutant strains of AB98041, the *hppD* mutant alleles containing different base mutations were amplified from the corresponding pUC-hppDAS variant plasmids. The amplified fragments were then linked to upstream and downstream arms by overlapping PCR to generate continuous products that contain the mutant alleles flanked by two arms. Similarly, to replace the *hppD* gene of AB98041 with the *hppD*-WS, the latter was amplified and inserted into the two arms. After sequencing, these constructs were inserted into pDM4 and transformed into *E. coli* β2155. Conjugation was performed between the β2155 strain and AB98041Δ*hppD*. The following procedures were identical to the construction of deletion mutants. The correct insertion of *hppD* mutant alleles and replacement of the *hppD* gene with *hppD*-WS in the AB98041 chromosome were all verified by PCR and sequenced for confirmation.

*hppD* deletion and replacement with *hppD*-WS in *A.s.s* strains 2013-8 and 2014-235 were also performed using the same approach.

### Purification and enzymatic characterization of wild-type and mutant enzymes

Colonies of *E. coli* BL21(DE3) expressing wild-type and mutant HppD enzymes were cultivated overnight in 5 ml LB medium containing kanamycin (50 μg/ml). The overnight cultures were inoculated into 100 ml of the same fresh medium until the OD_600_ reached ~0.6. IPTG was then added to a final concentration of 0.5 mM. After another 5 h of induction at 22 °C (16 °C and 12 h for HppD-AS), the cells were harvested from culture broth by centrifugation at 8,000 × *g* for 20 min at 4 °C, and then resuspended in 20 mM Tris-HCl buffer (pH 7.9) containing 500 mM NaCl, 10% glycerol, and 20 mM imidazole. The resuspended cells were disrupted on ice by sonication and the resulting lysate was centrifuged at 13,000 × *g* for 20 min at 4 °C to pellet insoluble material. The supernatant was filtered through a 0.22-μm filter (Merck Millipore, Germany) and passed over a Ni^2+^-NTA matrix (GE Healthcare, USA) according to the manufacturer’s instructions to purify His_6_-tagged enzymes. The column was washed with 20 mM Tris-HCl buffer (pH 7.9) containing 500 mM NaCl, 10% glycerol, and 40 mM imidazole. Proteins bound to the column were eluted with an imidazole gradient (80 to 500 mM) in buffer (20 mM Tris-HCl, 500 mM NaCl, and 10% glycerol, [pH 7.9]). All fractions were collected and analyzed by SDS-PAGE. Homogeneous peak fractions were pooled and concentrated using a centrifugal filter (Merck Millipore, Germany) equipped with a 10,000 MWCO membrane against 100 mM Tris-HCl buffer (pH 7.5) to simultaneously remove imidazole. Protein concentrations were determined by the Bradford assay, with bovine serum albumin as the standard.

HppD activity was measured as described previously (58) with some modifications. Briefly, the assay was performed in 100 μl of reaction buffer containing 100 mM Tris-HCl (pH 7.5), 1 mM Na-ascorbate, 250 μM ferrous sulfate, 1 mM 4-hydroxyphenylpyruvate (Sigma, USA), and 100 μg purified protein (depending on specific activity). The mixture was incubated for 30 min and then the reaction was terminated by the addition of acetonitrile (15% [v/v] final concentration). The precipitated proteins were removed by centrifugation at 16,000 × *g* at 4 °C for 5 min and the supernatant was subjected to HPLC analysis. An injection volume of 20 μl was used on an Agilent 1100 liquid chromatograph fitted with a C18 reverse phase column (4.6 × 250 mm; 5-μm particle size) and a DAD-UV detector. The mobile phase consisted of 90% (v/v) 10 mM acetic acid and 10% (v/v) methanol with a flow rate of 1 ml/min. HGA production was monitored at 290 nm and quantification was performed by measuring peak areas. The areas were transformed to nmol based on a standard curve obtained with commercially available HGA (Sigma, USA). One unit (U) of enzyme activity was defined as the amount of HGA (μmol) generated by per g enzyme per min. The optimal temperature of each enzyme was determined by measuring activity at temperatures ranging from 16 and 53 °C. Thermostability was determined by incubating the enzyme at 40 °C, and aliquots were then taken at different time points to determine residual activities at 22 °C as described previously herein.

### Homology Modeling

The possible effects of substitutions of critical residues on protein structural stability were evaluated by analyzing the three-dimensional structural model. A molecular model of the HppD-AS enzyme was generated by homology modeling using SWISS-MODEL (59).

The HppD of *Pseudomonas fluorescens* A32 (60) (PDB accession number 1CJX) that shared 51% identity with HppD-AS was used as the template. The PyMOL molecular graphics system was used to visualize and analyze the structure of wild-type HppD-AS and mutants.

## Acknowledgments

This work was supported by the National Natural Science Foundation of China (No. 31770052), the National Fund for Fostering Talents of Basic Sciences (J1103513), and Research (Innovative) Fund of Laboratory of Wuhan University. The funders had no role in study design, data collection, and interpretation, or the decision to submit the work for publication. We thank Dr. Tongyan Lu (Heilongjiang River Fisheries Research Institute, Chinese Academy of Fishery Sciences) for kindly providing *A.s.s* strains 2013-8 and 2014-235. There are no conflicts of interest to declare.

Yunqian Qiao and Xiangdong Chen contributed to the conception and design of the study. Yunqian Qiao, Jiao Wang, He Wang, Baozhong Chai, and Chufeng Rao performed the experiments and generated the data. Yunqian Qiao, Xiangdong Chen, and Shishen Du analyzed the data and wrote the paper.

